# Learning to count: determining the stoichiometry of bio-molecular complexes using fluorescence microscopy and statistical modelling

**DOI:** 10.1101/2020.07.23.217745

**Authors:** Sophia F. Mersmann, Emma Johns, Tracer Yong, Will A. McEwan, Leo C. James, Edward A.K. Cohen, Joe Grove

**Affiliations:** Department of Mathematics, Imperial College London, London, U.K; Institute of Immunity and Transplantation, Division of Infection and Immunity, University College London, London, U.K; Department of Clinical Neurosciences, UK Dementia Research Institute at the University of Cambridge, Cambridge, U.K; Medical Research Council, Laboratory of Molecular Biology, Cambridge, U.K

## Abstract

Cellular biology occurs through myriad interactions between diverse molecular components, many of which assemble in to specific complexes. Various techniques can provide a qualitative survey of which components are found in a given complex. However, quantitative analysis of the absolute number of molecules within a complex (known as stoichiometry) remains challenging. Here we provide a novel method that combines fluorescence microscopy and statistical modelling to derive accurate molecular counts. We have devised a system in which a given biomolecule is differentially labelled with spectrally distinct fluorescent dyes (label A or B), which are then mixed such that B-labelled molecules are vastly outnumbered by those with label A. Complexes, containing this component, are then simply scored as either being positive or negative for label B. The frequency of positive complexes is directly related to the stoichiometry of interaction and molecular counts can be inferred by statistical modelling. We demonstrate this method using complexes of Adenovirus particles and monoclonal antibodies, achieving counts that are in excellent agreement with previous estimates. Beyond virology, this approach is readily transferable to other experimental systems and, therefore, provides a powerful tool for quantitative molecular biology.

The statistical models used in our analysis are available here: https://github.com/sophiamersmann/molecular-counting, the raw data used for molecular counting can be found here: 10.5281/zenodo.3955142.

## Introduction

All cellular processes are driven by coordinated networks of dynamically interacting molecular partners. To successfully function, these components typically need to be assembled into specific complexes or clusters. For example, receptor signalling generally requires the co-location of various sensory, regulatory and stimulatory partners; the precise makeup of these assemblies can tune the nature of the signal and resultant physiological output. Molecular cell biology research has, thus far, largely focused on determining the identity of the components found in a given complex. However, it is becoming clear that the quantity of any given component is also vitally important. Quantifying the number of molecules, or stoichiometry, within an assembly can be used to understand its ultrastructure and, ultimately, to create complete molecular models of entire cellular structures, as has been demonstrated for HIV capsids and the neurological synapse (Briggs et al., 2004; Wilhelm et al., 2014).

There are various approaches to investigate the number of molecules within a given complex; for example calibrated biochemical analysis or cryo-electron microscopy (cryo-EM). However, such methods pose practical and/or technological barriers and, by their very nature, obscure heterogeneity within the sample. Single molecule localisation microscopy (SMLM) modalities, such as STORM and PALM, are potentially powerful techniques for counting (Lee et al., 2012; Coltharp et al., 2012; Fricke et al., 2015; Veatch et al., 2012; Stein et al., 2019). Nonetheless, these approaches typically require detailed *a priori* knowledge of the experimental system (for instance, a thorough evaluation of the ‘blinking’ characteristics of the fluorophores (Patel et al., 2019)) and/or a well understood reference sample with which to calibrate the measurement (Thevathasan et al., 2019). These steps need to be performed independently for each different experimental model and microscope set up; this creates a high barrier to implementation and can make these methods vulnerable to experimental variation and artefacts.

Here we outline an alternative, and potentially complementary, approach that combines (non-SMLM) fluorescent microscopy and statistical modelling to extract estimates of molecular numbers within a complex. The method requires differential binary labelling of a constituent (i.e. a protein of interest is labelled with fluorescent dye A or B); by appropriate mixing of the labelled components, any individual molecular complex can be simply scored as being positive or negative for a given label. The frequency of positive complexes is then analysed by statistical modelling to extract estimates of stoichiometry. This approach is simple and requires minimal calibration or *a priori* understanding of the experimental system.

We demonstrate the feasibility and accuracy of this approach by studying the stoichiometry of virus-antibody complexes. Adenovirus (AdV) is a non-enveloped DNA virus; its genome is enclosed within a proteinous shell, called a capsid (Nemerow et al., 2012). The major AdV capsid protein is hexon; this assembles into trimeric subunits, that are hexagonal in shape, which in turn arrange to form an icosahedron with 20 triangular faces (a molecular cartoon of the AdV particle is provided in Figure 3A). The AdV particle has 12 vertices, each of which are occupied by a pentameric subunit (formed of the penton base protein) and a receptor binding ‘spike’ (formed of the fibre protein).

Antibodies (Ab) that bind sites such as the spike can directly neutralise AdV by blocking receptor interactions, therefore preventing the virus particle from entering the cell. However, antibodies targeting the hexon protein (which makes up the majority of the capsid) do not necessarily interfere with the mechanics of virus entry (Fender et al., 1995; Wohlfart et al., 1985). Nonetheless, anti-hexon antibodies can prevent virus infection by recruiting the intracellular antibody-sensor TRIM21, which targets the virus for degradation and activates cell-intrinsic immune responses (Keeble et al., 2008; Mallery et al., 2010). Anti-hexon monoclonal antibody 9C12 inhibits AdV infection via this mechanism and is used as a prototypical system to investigate TRIM21. The stoichiometries of antibody and TRIM21 recruitment to incoming virus particles remain unclear and are likely to be a determinant of the nature of the resulting cellular response.

Previous studies, using alternative techniques, have provided estimates of the stoichiometry of AdV-9C12 complexes. Each AdV particle possesses 720 identical hexon proteins, each of these represents a potential binding site for 9C12. However, the hexon subunits are assembled as trimers, and are arranged in a specific geometry (as described above). Moreover, antibodie are bivalent, therefore each 9C12 molecule has two hexon binding pockets. Consequently, it is highly unlikely that each hexon molecule will be occupied by a single 9C12 molecule (i.e. 720 antibodies per virus particle). Analysis by cryo-EM, immuno-gold EM staining and calibrated fluorescence measurements suggest a true maximum stoichiometry of 100-200 antibodies per particle (McEwan et al., 2012; Varghese et al., 2004); this maximum is likely determined by the limits to antibody binding and packing enforced by the geometry of the particle. We have applied our counting method to the AdV-9C12 complex and generated estimates that are in good agreement with these previous studies, therefore validating our approach.

## System and Methods

### Strategy

We used differential binary labelling and statistical modelling to extract estimates of stoichiometry, our strategy is outlined in Figure 1; note that this approach can be generalised to apply to many other multi-component systems (i.e. how many protein x are found in assembly y?). The hidden truth is the number of antibodies bound to a virus particle; the Ab:virus stoichiometry is expected to increase with antibody concentration until it reaches a saturation point where the maximum number of Abs are bound (Figure 1A.).

**Fig. 1.**
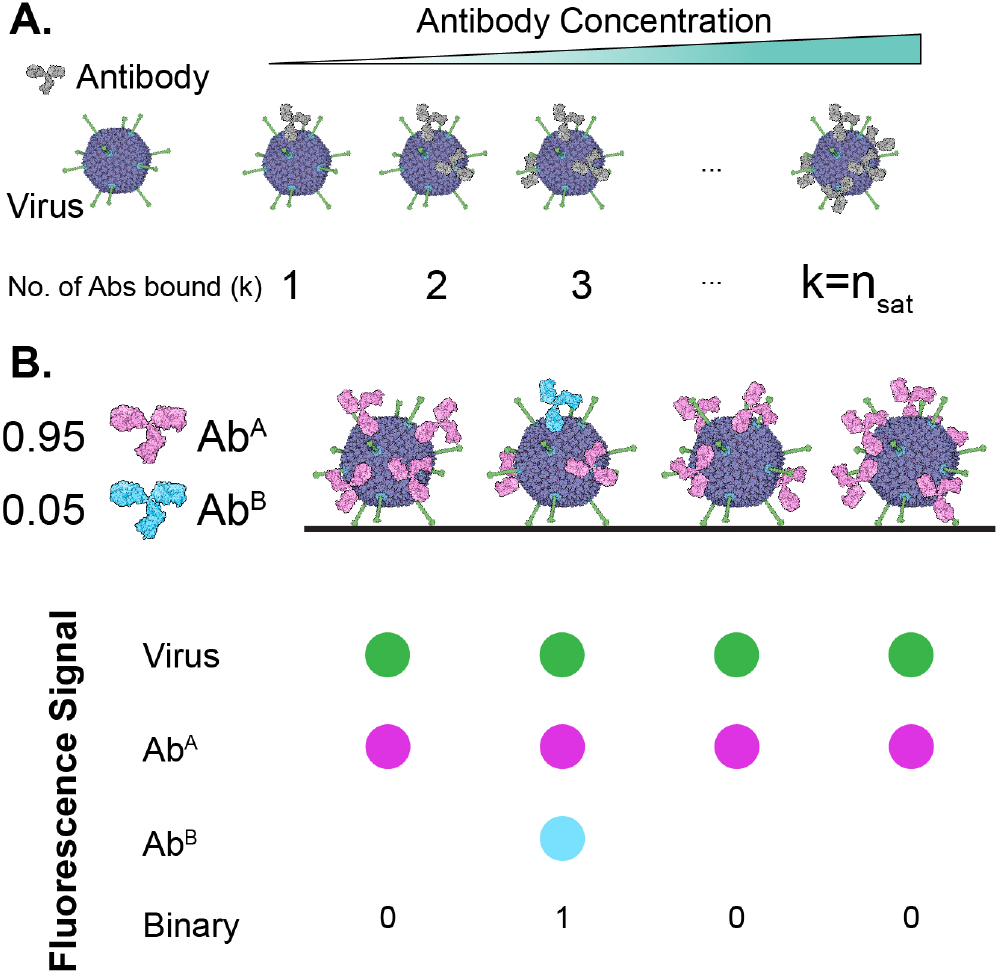
Binary Labelling of AdV-antibody Complexes. **A.** The ground truth: the number of antibody molecules (*k*) per virus particle increases with antibody concentration up to a saturation point (*k* = *n*_sat_). **B.** Extracting truth from data: AdV particles (labelled with a green fluorescent dye) are incubated with a defined mixture of two batches of antibody; one batch has received fluorescent label A (magenta), whilst the second has received label B (blue). When viewed by microscopy, every virus particle has received at least one molecule of AbA, whereas only a minority have received any AbB and can be scored in a binary fashion. Note, antibody molecules are not drawn to scale

In our method (Figure 1B), both components (virus and antibody) are fluorescently labelled, however, two separate batches of, the otherwise identical, antibody are given spectrally distinct dyes (resulting in Ab^A^ or Ab^B^). Mixing of the differentially labelled antibody batches at appropriate proportions results in only a minority of virus particles receiving a particular fluorescent dye (B in the example cartoon). Therefore, when imaged, we detect three distinct fluorescent signals: each virus particle can be identified by its reference dye (green in the cartoon example), every virus particle has also received antibodies with dye A (magenta), however, very few particles are positive for dye B (blue) and can be scored as positive or negative in a binary fashion. The frequency of virus particles that are positive for Ab^B^ is a function of the A:B mixing proportion and the stoichiometry of Ab:virus interaction; this relationship between the data and the hidden truth can be modelled.

Consider a single virus to be capable of binding *n*_sat_ antibodies at saturation. Under the assumption that antibodies bind to the same virus independently from each other, *K*, the total number of (labelled and unlabelled) antibodies bound to a virus, can be modelled as a binomial random variable

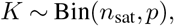

where *p* is the probability of an antibody binding to a particular binding site of the virus. If *n*_sat_ and *p* are known, then the expected number of antibodies bound to a single virus is simplyE[*K*] = *n*_sat_ ± *p.* However, in most cases *n*_sat_ is not known and *p* cannot be expressed easily since it depends not only on the antibody concentration used but also the composition of the virus particle and the geometry of interaction, which can be difficult to obtain. To extrapolate an antibody count it is, therefore, necessary to estimate both, *n*_sat_ and *p*.

As described above, our experimental design utilizes antibody labelled with spectrally distinct dyes allowing binary scoring of individual virus particles as positive if they interact with at least one Ab^B^ molecule (Figure 1). Here, we describe this state as being a Bernoulli random variable *S* that takes the value 1 if the virus is in the positive state, and 0 if it is in the negative state, i.e.

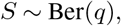

where *q* is the probability a virus interacts with at least one labelled antibody.

Since *q* = *P*(*S* = 1) = 1 – *P*(*S* = 0), we can derive a closed form for *q* by finding an expression for the probability of a virus not being complexed with any labelled antibody *P*(*S* = 0). To this end, we simply sum over all possible virus-antibody configurations under the constraint of all antibodies being unlabelled, i.e. a virus could bind zero, one, two, … up to *n*_sat_ unlabelled antibodies. The marginal probability of a virus being in a negative state is thus given by

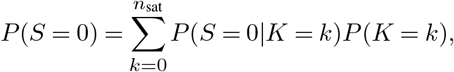

where *P*(*S* = 0|*K* = *k*) is the probability that given the virus binds to *k* antibodies, exactly zero of them are labelled. The conditional distribution of *S* given *K* = *k* is itself binomial, namely

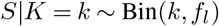

where *f_l_* is the proportion of antibodies that are fluorescently labelled. Therefore

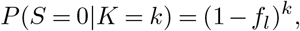

and combining with the probability mass function of *K*, namely

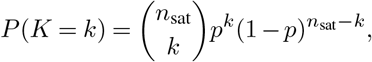

gives

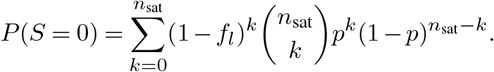

A closed form for *q* directly follows as

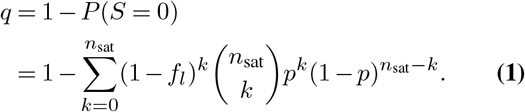

Note that here *P*(*S* = 0) is expressed in terms of *p* and *n*_sat_, and will hereafter be referred to as *P*(*S* = 0; ***θ***), where ***θ*** = (*p,n*_sat_).

Consider a single experiment (performed at a specific antibody concentration) to describe *V* viruses with states **s** = *s*_1_,…,*s_V_* where *V*_+_ of these states are positive, i.e. *V*_+_ viruses have been observed to interact with at least one labelled antibody. Assuming independence among viruses, the likelihood of ***θ*** is then

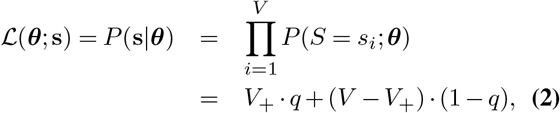

where *q* is as given in (1). We are then interested in the posterior distribution of ***θ*** given by the central result of Bayesian statistics,

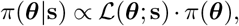

where *π*(***θ***) is a suitable prior for the unknown parameters ***θ***.

In most applications, more than one experiment is performed; consider *m* experiments to be conducted at varying antibody concentrations *c*_1_,…,*c_m_*, producing *m* sets of virus state observations 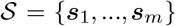. Parameter inference using the described single-experiment model would entail building *m* independent models, each estimating *p_j_* and *n*_sat,*j*_ for experiment *j*. While estimating concentration-specific antibody binding probabilities is desired, inferring multiple *n*_sat_ values is unintuitive since *n*_sat_ is fixed for a particular virus-antibody pair, i.e. *n*_sat_ should be common to all experiments regardless of the antibody concentration used. It is, therefore, preferable to develop a general model accounting for multiple experiments that estimates all *p*_1_,…,*p_m_* simultaneously while yielding only a single *n*_sat_ estimate.

Such a general model contains a single likelihood 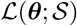, where ***θ*** = (*p*_1_,…,*p_m_*,*n*_sat_). This can be expressed as

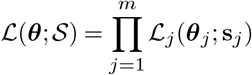

where 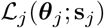 is the likelihood function for ***θ***_*j*_ = (*p_j_*, *n*_sat_) as defined in (2). Hence, *m* +1 unknown parameters are estimated; a probability *p_j_* specific to each concentration *c_j_* for *j* = 1,…,*m*, and crucially a single *n*_sat_ shared over all experiments. To sample from the posterior distributions Markov Chain Monte Carlo (MCMC) is used, specifically the Metropolis-Hasting algorithm using PyMC (Patil et al., 2010). No prior knowledge is incorporated by imposing a beta distribution Beta(1,1) on all *p_j_*, *j* = 1,…,*m*, and a uniform distribution with a sufficiently large upper bound, Uniform(0,1000), on *n*_sat_. The convergence of MCMC is checked by visual inspection of trace and autocorrelation plots for each parameter.

### Model verification via simulations

The proposed method makes use of experimental parameters including the proportion of antibodies labelled (*f_l_*) and the number of viruses sampled (*V*). Formally, these are not required to comply with specific bounds. However, certain configurations of an experimental set-up are not expected to yield data that lead to a sensible model. Labelling almost all antibodies (i.e. *f_l_* close to 1), for example, would result in a data set with low information content. To explore how different experimental design choices impact the model’s ability to reliably estimate parameters, we analysed simulated data that assumed a range of possible experimental settings.

We simulated a single experiment at a time and assumed the number of antibodies bound at saturation to be known. For this we used an upper limit estimate of AdV-9C12 stoichiometry that we previously derived from an alternative method, *n*_sat_ = 205 (McEwan et al., 2012). In each simulation, the binding probability *p* of an antibody is thus the only parameter estimated. We considered a range of possible experimental settings by varying the total number of viruses sampled in an experiment (from 100 to 4000) and the proportion of antibodies labelled (from 0.01 to 0.9). For an assumed true antibody binding probability *p* ∈ [0.1,0.99], data is simulated by drawing *V* virus states from *S* ~ Ber(*q*) with *q* as described in (1). The antibody binding probability was then blindly estimated using MCMC and convergence checked using the Gelman-Rubin statistic (Gelman and Rubin, 1992).

A total of 1331 simulations were carried out that assess the model’s ability to reliably estimate *p* in various experimental settings. As expected, high proportions of labelled antibodies produced data of low information content, reflected in the model’s inability to accurately estimate *p*, even if the number of viruses used in an experiment was high (Fig. 2A). By contrast, if *f_l_* is less than or equal to 0.1, *p* was estimated with low bias and low variance (Fig. 2A). Simulations also suggested that for low *f_l_*, as a rule of thumb, at least 500 viruses per experiment should be sampled (Fig. 2B-C). However, if the proportion of antibodies labelled is 0.1 (or higher), then the proposed model failed to produce a reasonable estimate of *p* for most underlying true values; higher number of viruses seemed to compensate this effect to some extent (Fig. 2D). In summary, these simulations put empirical bounds on experimental parameters and show that, if compliant, the method yields sensible estimates.

**Fig. 2.**
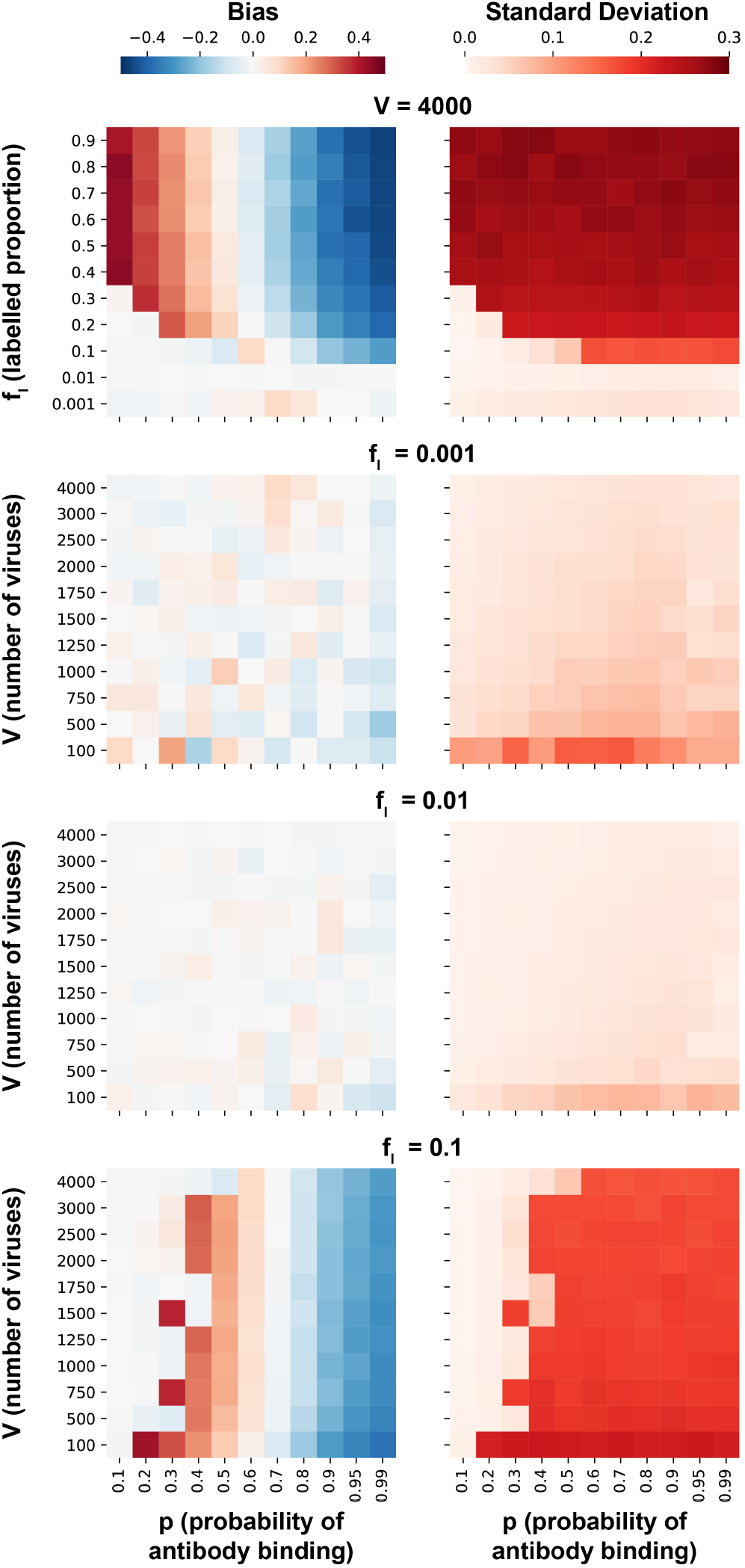
Bias and standard deviation landscapes of the posterior distributions of antibody binding probabilities estimated from simulated data that explore a range of experimental settings. Here, bias is the mean of the posterior distribution minus the true probability, while standard deviation refers to the standard deviation of the posterior distribution. **A.** Bias and standard deviation as a function of the true binding probability, *p*, and the proportion of antibodies labelled, *f_l_*. The number of viruses used in an experiment, *V*, is fixed at 4000. **B-D.** For a fixed value of *f_l_* (0.001, 0.01, and 0.1, respectively), bias and standard deviation are shown as a function of *p* and *V*.

**Fig. 3.**
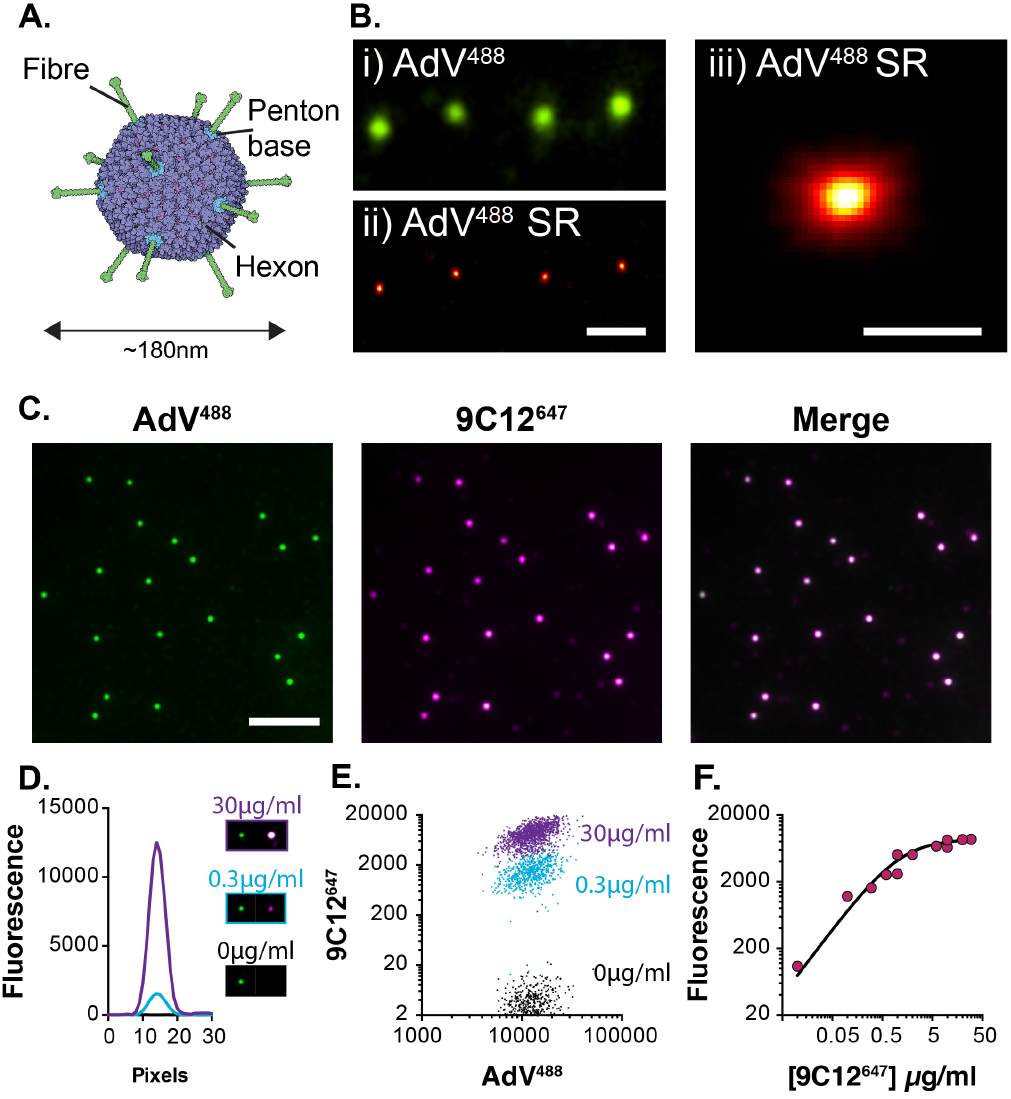
Experimental measurements of AdV-antibody complexes. **A.** A molecular cartoon of an AdV particle; Ab 9C12 targets the hexon protein. **B.** Immobilised AdV^488^ particles were imaged by TIRF-M (i) and SRRF (ii), iii provides an enlarged image of a super-resolved particle. Scale bars 1μm (i & ii) and 200nm (iii). **C.** A representative field of AdV^488^ particles incubated with 1μg/ml 9C12^647^, the merge image displays an overlay of both channels. Scale bar 5μm. **D.** 9C12^647^ fluorescent signals associated with AdV particles incubated with 30, 0.3 and 0μg/ml antibody.**E.** Fluorescent measurements of populations of AdV particles. **F.** Mean 9C12^647^ signals upon increasing concentration of antibody

## Implementation

### Experimental Setup

Successful implementation of our strategy requires sensitive and unambiguous measurements of individual virus-antibody complexes. We achieved this by immobilising AdV particles onto coverslips for analysis by total internal reflection fluorescence microscopy (TIRF-M; a detailed description of experimental methodology is provided in the supplementary information). Prior to immobilisation, purified AdV particles were directly labelled with Alexa Fluor 488; when observed by TIRF-M (Figure 3Bi) they appear as monodisperse diffraction limited spots (the particle being 180nm in diameter; Figure 3A). To further verify that each object is a single virus particle we used SRRF analysis (super-resolution by radial fluctuations (Gustafsson et al., 2016)); for each object we resolved a single maxima of fluorescence that was ~200nm in diameter (Figure 3Bii and iii), consistent with the ultrastructure of AdV particles (Figure 3A).

Immobilised AdV particles were incubated with monoclonal antibody 9C12 conjugated to Alexa Fluor 647 dye, and imaged by TIRF-M. Each AdV particle was positive for antibody, indicating the assembly of virus-antibody complexes (Figure 3C). Moreover, individual AdV-9C12647 complexes could be analysed in a quantitative manner over a >100 fold range in antibody concentration (Figure 3D). We automated this process to allow measurements of whole populations of virus particles at varying concentrations of antibody (Figure 3E & F). 9C12^647^ signal intensity was proportional to antibody concentration and reached a plateau at high values; this indicates increasing virus-antibody stoichiometries up to a saturation point at which maximum antibody binding is achieved (as outlined in Figure 1A). Moreover, the populations of virus particles were quite homogenous in 9C12 signal; this suggests a relative uniformity of assembly. In conclusion, we were able to quantitatively analyse the assembly of individual virus-antibody complexes.

To achieve differential labelling a second batch of 9C12 was directly conjugated with biotin; this served as the ‘B’ batch of antibody to be mixed with the 9C12^647^ ‘A’ batch (Figure 1). It may be possible that conjugation with either biotin or Alexa Fluor 647 has unexpected detrimental effects on antibody binding; therefore, to have confidence in our binary labelling system we needed to demonstrate fair competition between our differentially labelled antibody batches. To test this we incubated immobilised AdV particles with a high concentration of antibody (20μg/ml) composed of different proportions of 9C12^647^ or 9C12^Biotin^ (e.g. 0.75:0.25, 0.5:0.5). We then monitored the 9C12^647^ fluorescence signal under each condition. As the proportion of 9C12^647^ dropped we measured a stepwise reduction in fluorescence signal (Figure 4A). If both batches of antibody possess equivalent binding to AdV we would expect a linear relationship between the proportion 9C12^647^ and fluorescent signal. Indeed, when normalised for units, we observed a near perfect linear relationship (Figure 4B, slope = 0.99, R^2^=0.97); indicating a fair competition in binding between our A and B batches of antibody.

**Fig. 4.**
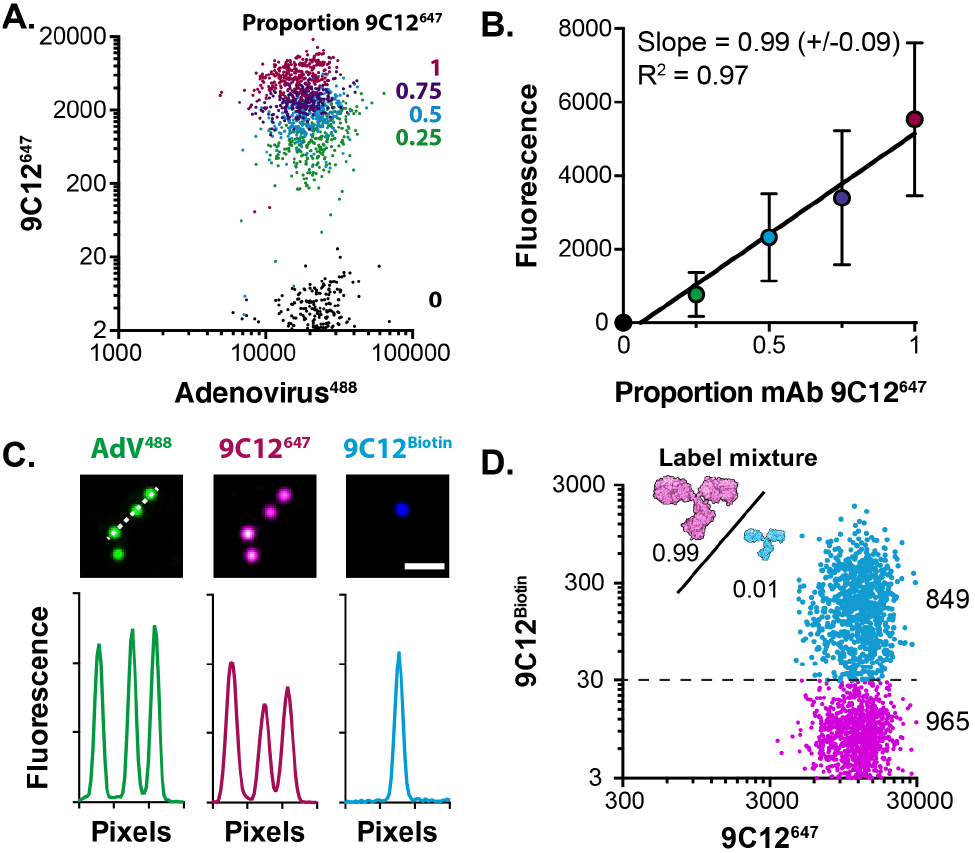
Binary labelling of AdV-antibody complexes. **A.** AdV particles were incubated with mixtures of mAb 9C12 labelled with either Alexa Fluor 647 or Biotin, to a final concentration of 20μg/ml. The scatter plot displays AdV^488^ and 9C12^647^ fluorescent signals for populations of AdV particles labelled with decreasing proportions of 9C12^647^, as stated in the legend. **B.** The mean 9C12^647^ signal is linearly proportional to the antibody mixture, this indicates equivalent binding by either batch of antibody. **C.** AdV^488^ particles (green) were labelled with 5μg/ml 9C12^647^ (magenta) spiked with 1% (0.01) 9C12^Biotin^ (blue); only a minority of particles received any 9C12^Biotin^. **D.** Scatter plot displaying a population of AdV particles (labelled as in C), which can be scored as being positive or negative for 9C12^Biotin^ (as annotated on the plot)

As depicted in Figure 1B, our approach requires detection of single antibody molecules (of B batch) within individual virus-antibody complexes. To explore this we incubated immobilised AdV particles with 5μg/ml 9C12^647^ spiked with 1% 9C12^Biotin^ (i.e. f_l_ 0.99:0.01). Molecules of 9C12^Biotin^ were detected using streptavidin (an ultra-high affinity biotin binding protein) conjugated to a quantum dot (QDot^655^). The photostability of quantum dots permits signal accumulation over prolonged exposure times (Resch-Genger et al., 2008; Algar et al., 2011), therefore increasing the sensitivity of detection. Analysis by TIRF-M revealed that whilst every particle was positive for 9C12^647^ only a subset possessed 9C12^Biotin^-QDot^655^ signal (Figure 4C); this suggests a population of AdV particles receiving one, or very few, 9C12^Biotin^ antibody molecules (as outline in Figure 1B). Automated analysis revealed well-separated populations of AdV particles that were positive or negative for 9C12^Biotin^ (Figure 4D). Note that all particles were positive for 9C12^647^. These data are consistent with the assembly of virus-antibody complexes in which the vast majority of antibody molecules are from batch A (9C12^647^), but a subset of complexes contain ~1 molecules of batch B antibody (9C12^Biotin^); the proportion of 9C12^Biotin^ positive particles will serve as the output data for statistical modelling.

### Results

We have demonstrated quantitative analysis of 9C12 interaction with individual Adv particles (Figure 3); we have confirmed that differential labelling of antibody does not bias binding (Figure 4A & B); and that we could detect single molecules of 9C12^Biotin^ allowing discrimination of positive and negative AdV-9C12 complexes (Figure 4C & D). Therefore, we have satisfied the necessary technical requirements to implement the strategy outlined in Figure 1.

We proceeded with a series of experiments to generate data for statistical modelling and stoichiometric estimates. To achieve this we performed four independent overlapping titrations over a >100 fold range in antibody concentration (0.15-20μg/ml). Guided by the simulations in Figure 2, we explored a range of 9C12^Biotin^ proportions, from 0.007-0.07 (0.7-7%), and, where possible, collected >500 particles per sample (the average number of particles collected was >1500). The proportion of positive particles was assessed for each sample, details of data analysis are provided in the supplementary information.

Supplementary Figure 1 provides representative data: scatter plots display AdV^488^, 9C12^647^ and 9C12^Biotin^ intensities for control samples (treated with unmixed 9C12^Biotin^ or 9C12^647^) and four representative test samples incubated with a range of antibody concentration, containing 0.007 9C12^Biotin^. Bar charts provide summary statistics for each channel: the particles have uniform AdV^488^ reference signal, whereas the 9C12^647^ fluorescence decreases with antibody concentration; likewise, the proportion of 9C12^Biotin^ positive particles decreases with concentration. This data are consistent with the expected concentration-dependent stepwise reduction in virus-antibody stoichiometry.

The four experiments generated 24 sets of binary scores (Supplementary Figure 1 and Table 1) which were integrated in to our statistical model, as outlined in the methods. This generated posterior distributions of p (probability of an antibody binding site being occupied) for each sample (Supplementary Figure 2A) and an estimated nsat (the maximum number of antibodies that can bind per particle) of 133 (maximum a posteriori (MAP) estimate; CI (credible interval) [119,162]; Supplementary Figure 2B). Absolute antibody numbers for each sample can be derived by multiplying the MAP estimate of each posterior distribution (for p) by nsat (Supplementary Figure 2C). Figure 5A displays the mean number of bound antibody at increasing 9C12 concentrations; AdV-9C12 stoichiometries range from 29 to 115 across the titration of antibody. For any given sample, alongside the proportion of particles that are positive for 9C12^Biotin^, the experimental setup provides fluorescent intensity values of 9C12^647^ in each AdV-Ab complex (Supplementary Figure 1); this provides an internal reference for AdV-9C12 interaction, therefore, the stoichiometric estimates should correlate with their experimentally-matched fluorescent intensity values. Figure 5B demonstrates a near perfect linear correlation between stoichiometry and fluorescence intensities for an example experiment; this would suggest that our statistical modelling faithfully reports the relationship between antibody concentration and AdV binding occupancy.

**Fig. 5.**
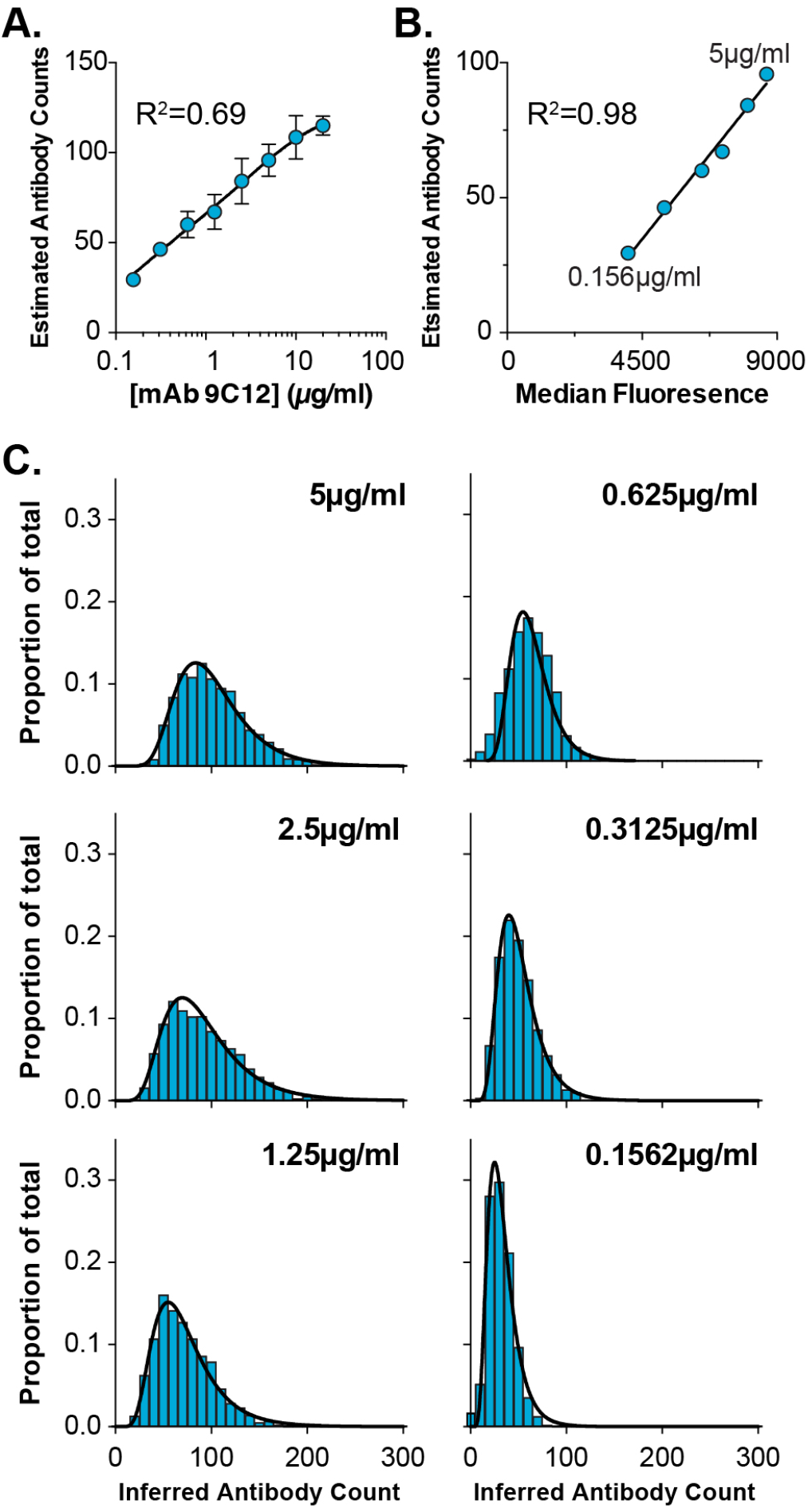
Multiple titrations of 9C12^647^ (containing 0.007-0.07 9C12^Biotin^) were performed to allow statistical modelling of AdV-9C12 interaction stoichiometries. **A.** Mean stoichiometric estimates at increasing concentrations of 9C12, error bars indicate standard error of the mean; data fitted with a binding curve, R^2^ = 0.69 (Graphpad, Prism). **B.** Stoichiometric estimates were compared to 9C12^647^ fluorescent values from an individual experiment; data fitted with a linear regression, R^2^ = 0.98 (Graphpad, Prism). **C.** Stoichiometric estimates were used to calibrate fluorescent intensity values allowing inference of heterogeneity. Histograms display the frequency of inferred antibody counts as a proportion of total particles. The frequency data were fitted with a log-normal distribution, R^2^ values were all >0.99 (Graphpad, Prism).

The stoichiometric estimates generated by our method derive from an ensemble measurment and, therefore, represent the antibody interactions of the average virus particle; this obscures heterogeneity within the population. However, given the excellent agreement between the estimated antibody counts and 9C12^647^ intensity values (Figure 5B) we used the stoichiometric estimates to calibrate the 9C12^647^ signals, therefore allowing us to infer population heterogeneity. To achieve this, for any given sample we matched the median 9C12^647^ fluorescence intensity to its associated stoichiometic estimate (generated by the model); extrapolating from this we then converted the 9C12^647^ fluorescence values to inferred antibody counts for individual virus particles. Figure 5C provides histograms and frequency distributions of inferred antibody counts for a range of concentrations. Being slightly skewed to the right the frequency data was best fitted using a log-normal distribution, this is particularly apparent at higher antibody concentrations. This would suggest that a significant proportion of virus particles are binding more antibodies than the average particle. However, no particles bound greater than 230 antibody molecules. We provide an interpretation of these observations in the discussion.

In summary we have used statistical modelling to derive stoichiometric estimates of AdV-9C12 complexes. This suggests that the most probable antibody binding maximum is 133 molecules (95% CI 119-162). However, using stoichiometric estimates to calibrate fluorescent data revealed population heterogeneity with a small proportion of virus particles binding ^200 antibody molecules. Notably, these values are in excellent agreement with previous estimates that we, and others, have derived using alternative methods (McEwan et al., 2012; Varghese et al., 2004).

## Discussion

Various methods permit the investigation of molecular stoichiometries within biological assemblies but technical limitations often make it difficult to obtain reliable estimates. For example SMLM, by its very nature, identifies single molecules and, if properly calibrated, can deliver accurate stoichiometries; however, successful counting by SMLM requires a very detailed understanding of the photochemical behaviour of the chosen fluorescent dyes. Here we outline a robust, and relatively facile, experimental framework for extracting accurate molecular counts using (non-SMLM) fluorescent microscopy and statistical modelling

Labelling of a component of interest with spectrally distinct fluorescent dyes (label A or B), and mixing them at defined ratios such that B-labelled molecules are in the minority, allowed assembled complexes to be simply identified as being either positive or negative. The frequency of positive complexes was then related to the underlying stoichiometry of interaction through statistical modelling.

By creating a scheme in which complexes need only be qualitatively scored for a particular label, our method negates the necessity for carefully calibrated measurements and an *a priori* understanding of the system. The only requirement is that the chosen label is clearly discerned from background; this is easily achievable with bright/stable fluorescent dyes and relatively inexpensive cameras; in this case we used quantum dots for sensitive detection, but many standard fluorescent dyes should also suffice.

An obvious limitation is that our method relies on an ensemble measurement and obscures heterogeneity within the population of complexes. However, this information remains accessible via the A-label fluorescent signals measured from each complex. Consequently, the ensemble-based stoichiometric estimates of the average complex can be used to calibrate these fluorescent signals and infer approximate molecular counts for individual complexes, therefore, restoring heterogeneity.

We were able to make robust measurements of AdV-9C12 interactions by fluorescence microscopy and successfully implemented the differential labelling strategy. We performed multiple independent measurements at various antibody concentrations to derive molecular counts for AdV-9C12 complexes. Our analysis estimated that the average AdV particle interacts with a maximum of 133 9C12 antibody molecules. Moreover, through examination of population heterogeneity we revealed that a small proportion of AdV particles may bind up to 230 antibody molecules.

These data are can be reconciled with a molecular model of AdV-9C12 complexes. AdV particles possess 720 identical hexon subunits, each providing a potential binding site for 9C12, however, previous estimates indicate an absolute maximum of 240 antibody molecules per virion. This would suggest that particle geometry places packing constraints on the arrangement of antibody molecules. Cryo-EM analyses indicates a complex and heterogeneous interaction network in which particle geometry creates potential antibody clashes and, therefore, prevents binding to every site simultaneously (Varghese et al., 2004). Whilst there are consistently five 9C12 molecules at each of the twelve vertices of the AdV particle, additional antibody interactions occur through heterogeneous packing across the surface; the pattern of which is likely dictated by the random order in which binding sites become occupied on any given virus particle. As a consequence with optimal antibody packing there is likely to be an absolute maximum binding occupancy of ~240 molecules. Our data indicate that the average particle binds fewer molecules (133) than the threshold dictated by purely geometric limitations. This would suggest that the majority of particles do not achieve optimal antibody packing and saturate at lower occupancies. This model also offers explanation to our observation that no particles bind greater than ~230 molecules (Figure 5C).

This interpretation of our data exposes another potential flaw in our approach. Our modelling strategy assumes that components bind independently, but in this test case the geometry of AdV particles create clashes between adjacent 9C12 molecules such that complete saturation is not possible. Consequently, antibody binding events can be influenced by prior antibody interactions and, therefore, are not independent. Although this may have introduced inaccuracies in our estimates, the molecular counts derived from our approach are in good agreement with previous values. Moreover, we maintain that the assumption of independent binding is appropriate for a generalisable method that can be applied to other biological assemblies. For example, within virology, we intend on extending this method to investigate the molecular composition and antibody-mediated neutralisation of enveloped viruses such as human immunodeficiency virus, hepatitis C virus and SARS-coronavirus-2. Beyond the confines of virology, our method could be applied to a variety of other biological assemblies of various scales, for example: bacteria; purified cellular organelles; cellular vesicles, such as exosomes; and supramolecular complexes such as ribosomes or inflammasomes.

In conclusion, we have developed a novel and robust method for counting components within biomolecular complexes. This approach has provided accurate counts for a previously characterised system and could be applied in a variety of other contexts. Moreover, we expect this system could be integrated with other complementary methods to enhance quantitative analysis; for example differential labelling and statistical modelling may provide a means of internally calibrating SMLM-based counting schemes.

## ACKNOWLEDGEMENTS

We would like to thank Ricardo Henriques for initiating some important conversations and providing technical assistance during image analysis. We would like to thanks Niall Adams for his help in supervising SFM during the development of the statistical methodology. Molecular cartoon images were taken from the Wellcome Collection digital image library. Both images are accredited David S. Goodsell, RCSB Protein Data Bank, and are used under an Attribution 4.0 International (CC BY 4.0) licence. JG is supported by a Sir Henry Dale Fellowship from the Wellcome Trust and Royal Society (107653/Z/15/Z).

