## Supplementary Information for "Learning to count: determining the stoichiometry of bio-molecular complexes using fluorescence microscopy and statistical modelling"

Mersmann et. al. 2020

### Supplementary Methods

#### *Virus preparations*

Replication-deficient human adenovirus-5 stocks were generated in a transcomplementation 293F cell line for 72 hours and purified by two rounds of CsCl centrifugation. Purified particles were conjugated to Alexa Fluor 488 using a protein labelling kit (Thermo Fisher Scientific).

#### *Antibody production and labelling*

9C12 anti-hexon monoclonal antibody was purified from hybridoma supernatants on a protein G column (GE Healthcare). Protein labelling kits were used to conjugate 'A' batch antibody to Alexa Fluor 647 (Thermo Fisher Scientific) and 'B' batch antibody to biotin (Merck Life Sciences). The degree of biotinylation was determined using a fluorescence biotin quantitation kit (Thermo Fisher Scientific); this was important for achieving accurate counts.

#### *Sample preparation*

25mm coverslips received two 2 minute washes in each of the following $ddH_20$, ethanol and methanol. They were then treated with 2% 3-amino-propyltriethoxysilane (Merck Life Sciences), diluted in acetone, for 5 minutes before being rinsed twice in $ddH_2O$. Each slide was mounted in an Attofluor chamber (Thermo Fisher Scientific) to which AdV, diluted in PBS, was then added. The samples were incubated in the dark overnight in a humidified box at 4°C, with gentle shaking.

The following day, the samples were rinsed in PBS and then blocked for 30 minutes in 1% BSA (diluted in PBS and 0.2µm filtered). Following two PBS rinses the samples were then incubated for one hour with 9C12 antibody (at various concentrations and combinations of A and B batches, as stated in the text). Unbound antibody was washed away by two sequential rinses in PBS, the samples were then fixed for 15 minutes in 4% EM-grade formaldehyde (Thermo Fisher Scientific), followed by two further PBS rinses. To detect biotinylated antibody, samples were incubated for 15 minutes with 2nM QDot 655 streptavidin conjugates, followed by two final PBS rinses. Samples were gently agitated on a rocking platform throughout the staining procedure. PBS was used as diluent at all stages.

We found air-drying to be very detrimental to sample preparation and therefore minimised the opportunity for this to occur throughout the entire sample preparation process. For example we used two P1000 pipettes when rinsing coverslips; one to aspirate, one to add buffer as soon as possible.

***Microscopy***
Samples were imaged on a Nikon Ti inverted microscope (Nikon Instruments), through a 100X oil objective, with 405nm (QDot), 488nm (Alexa Fluor 488) and 635nm (Alexa Fluor 647) laser illumination in TIRF mode. Images were captured using a 1024x1024 region of interest on a ORCA-Flash 4.0 sCMOS camera (Hamamatsu).

***Data Analysis***
Images were processed in ImageJ using a custom image analysis macro, which includes the channel align tool from the NanoJ package (Ricardo Henriques, https://github.com/HenriquesLab). The analysis pipeline outputs fluorescent intensity values for each particle in all three channels; particles were scored positive for $9C12^{Biotin}$ (B label) if they had a signal >30 (2X standard deviations above background). Scoring was manually verified for a random selection of images. The raw imaging data used to derive molecular counts, and the ImageJ anlaysis macro, are provided here: 10.5281/zenodo.3955142.

| Experiment | 1 | | 2 | | 3 | | 4 | |
|---|---|---|---|---|---|---|---|---|
| Proportion 9C12$^{Biotin}$ ($F_l$) | 0.0750 | | 0.0075 | | 0.0225 | | 0.0225 | |
| | n | Proportion +ve | n | Proportion +ve | n | Proportion +ve | n | Proportion +ve |
| 20 | 1190 | 1 | 3143 | 0.545020681 | 427 | 0.941451991 | | |
| 10 | 648 | 1 | 3070 | 0.46970684 | 839 | 0.94398093 | | |
| 5 | 900 | 1 | 1976 | 0.430161943 | 1072 | 0.906716418 | 1160 | 0.912068966 |
| 2.5 | 862 | 1 | 1632 | 0.330882353 | 1414 | 0.859971711 | 2541 | 0.866587957 |
| 1.25 | 799 | 0.998748436 | 2258 | 0.291851196 | 1003 | 0.77666999 | 1332 | 0.795045045 |
| 0.625 | 1254 | 0.997607656 | 3117 | 0.306705165 | 725 | 0.728275862 | 1840 | 0.70326087 |
| 0.3125 | | | | | | | 1793 | 0.658672616 |
| 0.15625 | | | | | | | 4237 | 0.47911258 |

**Supplementary Table 1. Outputs used for statistical modelling of virus-antibody stoichiometries.** We performed four independent overlapping antibody titrations at varying mixing proportions of 9C12$^{647}$ and 9C12$^{Biotin}$. The proportion of 9C12$^{Biotin}$ is stated for each experiment. The table provides the number of virus-antibody complexes that were analyzed (n) and the proportion of positive complexes, for each antibody concentration.

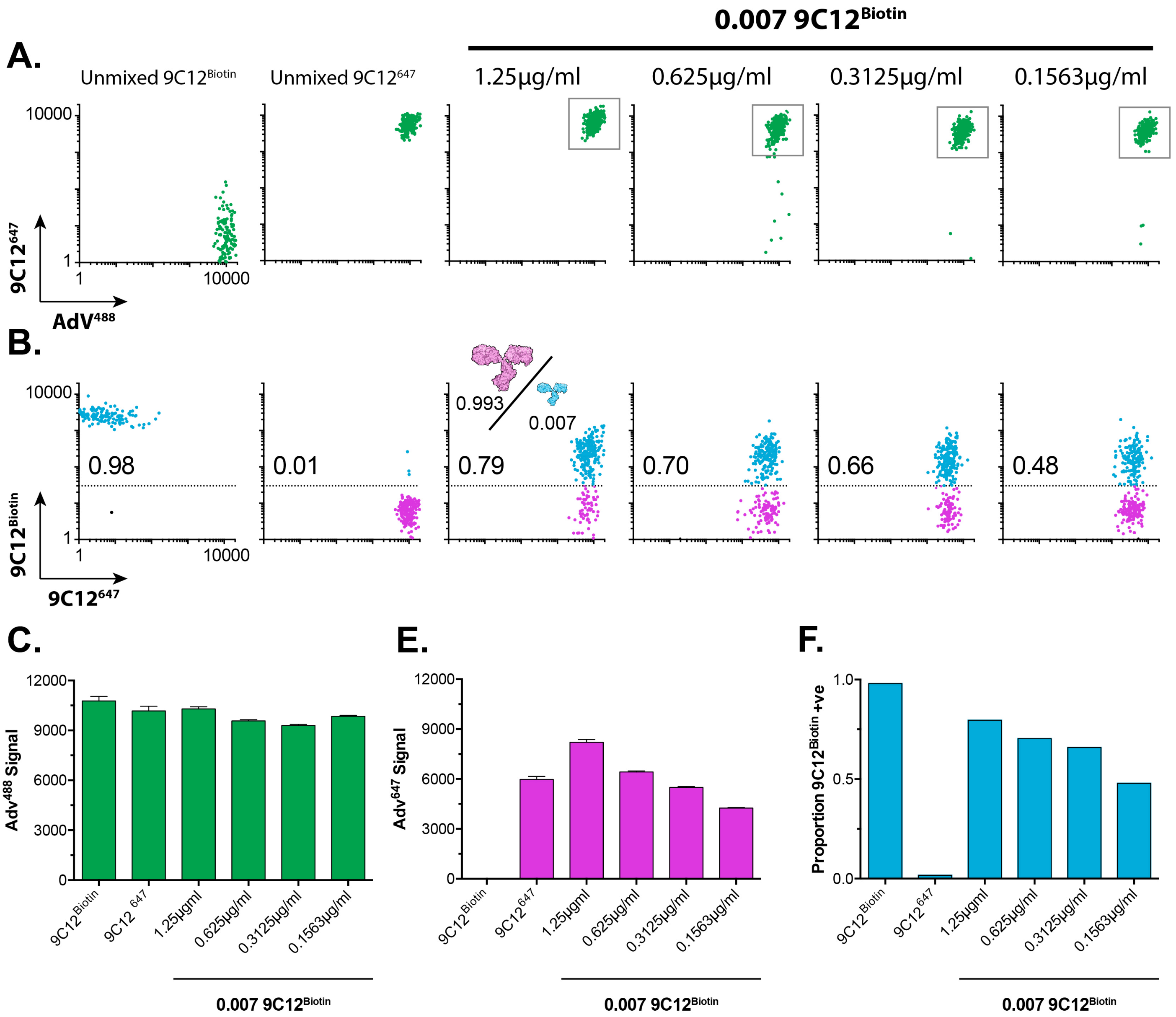

**Supplementary Figure 1. Quantification of differentially labelled virus-antibody complexes.**
**A.** Scatter plots displaying AdV$^{488}$ and 9C12$^{647}$ signals, for clarity 300 representative particles are shown. Control samples labelled with 100% 9C12$^{Biotin}$ or 9C12$^{647}$ (0.625µg/ml) are on the left, additional plots display particles labelled with stated concentrations of 9C12$^{647}$ spiked with 0.007 9C12$^{Biotin}$. Downstream analysis of 9C12$^{Biotin}$ was performed only on particles with 9C12$^{647}$ signal, as illustrated by the grey boxes. **C.** Scatter plots displaying 9C12$^{647}$ and 9C12$^{Biotin}$ signals from the same particles as shown in B. 9C12$^{Biotin}$ positive particles (signal ≥30, dotted line) are color coded in blue, the proportion of positive particles is annotated for each sample. **D.** The mean AdV$^{488}$ fluorescent intensity signals for each sample; ~2000 particles were analysed in each condition. **E.** The mean 9C12$^{647}$ fluorescent intensity signals for each sample. **F.** The proportion of 9C12$^{Biotin}$ positive particles for each sample. Error bars indicate standard error of the mean.

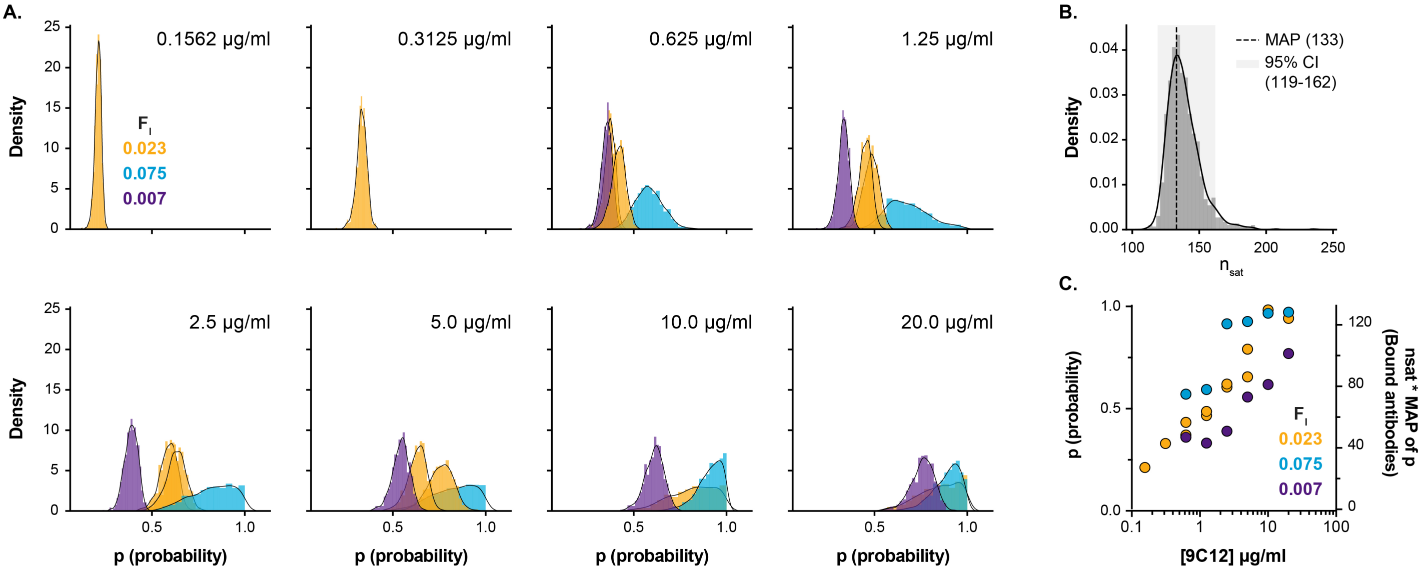

**Supplementary Figure 2. Posterior distributions and estimated parameters of the proposed statistical model. A.** Posteriors of antibody binding probabilities (p) for all measurements listed in Supplementary Table 1, grouped by the antibody concentration used in an experiment. Colours indicate the proportion of antibodies that are fluorescently labelled in an experiment ($f_l$). **B.** Posterior distribution of $n_{sat}$, the number of antibodies bound to a virus at saturation. The maximum a posteriori (MAP) estimate of $n_{sat}$ is 133 and lies within a 95% credible interval of [119, 162]. **C.** Scatter plot displaying MAP estimates of all binding probabilities inferred from the posterior distributions shown in A, plotted against antibody concentration. The axis on the right additionally shows the expected number of bound antibodies ($n_{sat}$ * MAP of p).
